# Multi-omics Data Integration by Generative Adversarial Network

**DOI:** 10.1101/2021.03.13.435251

**Authors:** Khandakar Tanvir Ahmed, Jiao Sun, Sze Cheng, Jeongsik Yong, Wei Zhang

## Abstract

Accurate disease phenotype prediction plays an important role in the treatment of heterogeneous diseases like cancer in the era of precision medicine. With the advent of high throughput technologies, more comprehensive multi-omics data is now available that can effectively link the genotype to phenotype. However, the interactive relation of multi-omics datasets makes it particularly challenging to incorporate different biological layers to discover the coherent biological signatures and predict phenotypic outcomes. In this study, we introduce omicsGAN, a generative adversarial network (GAN) model to integrate two omics data and their interaction network. The model captures information from the interaction network as well as the two omics datasets and fuse them to generate synthetic data with better predictive signals. Large-scale experiments on The Cancer Genome Atlas (TCGA) breast cancer, lung cancer, and ovarian cancer datasets validate that (1) the model can effectively integrate two omics data (e.g., mRNA and microRNA expression data) and their interaction network (e.g., microRNA-mRNA interaction network). The synthetic omics data generated by the proposed model has a better performance on cancer outcome classification and patients survival prediction compared to original omics datasets. (2) The integrity of the interaction network plays a vital role in the generation of synthetic data with higher predictive quality. Using a random interaction network does not allow the framework to learn meaningful information from the omics datasets; therefore, results in synthetic data with weaker predictive signals.

## Introduction

Complex diseases such as cancer are highly heterogeneous with different subtypes leading to varying clinical outcomes including prognosis, response to treatment, and chances of recurrence and metastasis (1–3). Disease phenotype prediction has been the subject of interest to clinicians and patients for many decades. The recent developments in high throughput sequencing technologies are capable of measuring molecular activities in cells and allow researchers to obtain multiomics data with sufficient quality and yield (4). It has revolutionized medical and biological research by offering a more comprehensive view of the underlying biological process of disease and identify accurate molecular signatures for characterizing or predicting disease phenotypes. Analysis of multiomics data along with clinical information of patients can help bridging the gap between genotype and phenotype by exploring the flow of information within different omics layers (5). These omics layers provide non-redundant predictive signals for predicting therapeutic response. Removing one of them in a prediction system will lead to performance degradation (6). Therefore, multi-omics data may provide a complementary set of information to understand the molecular basis of diseases. Predicting the phenotype using the multiomics data as independent sets of features will fall short in characterizing the prediction.

Disease phenotypes depend on molecular profiles and interplay at genomic, epigenomic, transcriptomic, proteomic, and metabolomic levels (5) which are interconnected with each other through complex networks (7). For instance, microRNA (miRNA) regulates mRNA expression by complementarily binding to recognition sequences in the 3’ untranslated region of their target mRNAs leading to mRNA degradation and/or mRNA translation inhibition (8). The abundance of a particular miRNA does not illustrate the full picture without knowing which mRNAs get inhibited by that miRNA; because miRNA does not directly influence the phenotype; rather, regulates the mRNA translation into protein that subsequently determines the phenotype. Moreover, mRNA can be regulated by other modulators like RNA binding protein (RBP) (9). RBPs bind RNA through globular RNA-binding domains (RBDs) and alter the expression of the bound RNAs (10). RNA-RBP interaction obtained from crosslinking and immunoprecipitation-based CLIP-Seq can also be applied to characterize the relation between omics data. Hence, integrating the interaction network into multiomics data analysis will capture the regulatory effect and establish a better correlation with the phenotype.

Several advanced multi-omics data integration frameworks have been proposed in the last five years (11–14). However, few approaches link different omics profiles using molecular interaction (15). Most of them ignore the relations across different biological layers in their analysis. The power of high throughput technologies cannot be fully utilized unless the multi-omics data with its intermodal relations are considered in studies.

In recent years, generative adversarial networks (GAN) (16) has gained popularity in solving problems within the scope of computational biology. GANs take random noise or pre-defined data as input and generate plausible synthetic data similar to a real dataset by imitating the distribution of the real data. There are several studies that use GAN based algorithms to generate data from single or multiple omics datasets. (17) used GAN for better biomarkers identification by generating a reconstructed functional interaction network from multi-omics datasets. (18) integrated diverse single-cell RNAseq (scRNA-seq) datasets from different labs and experimental protocols to simulate realistic scRNA-seq data that covers the full cell type diversity. (19) on the other hand used GAN to generate gene expression from bulk RNA-seq datasets. Recently, (20) proposed framework to eliminate batch effect from scRNA-seq data using GAN. GANs can learn non-linear relationships between features of omics data during training that can be used later for additional insight (18). It can handle missing data and also promising for missing value imputation because of its capability of learning and imitating any distribution of data (21). Based on its property of imitating distribution, we can design a GAN with one omics data from one distribution as input to the generator and another omics data with different distribution as real dataset in the discriminator to generate a synthetic data retaining information from both omics datasets.

In this study, we propose a biologically-motivated deep learning-based model, omicsGAN, to predict disease phenotype by integrating two omics data and the interaction between them (e.g., mRNA expression, miRNA expression, and miRNA-mRNA interaction network). The proposed model introduces a generative adversarial method to generate a new enriched feature set for each omics data combining information from the other omics dataset and the interaction network resulting in a better prediction. Experimental results verify that our proposed framework generates datasets with stronger molecular signatures to better understand the biological mechanism that leads to the disease state and improve disease outcome prediction compared to the biological features derived from single or concatenated omics data.

## Methods

In this section, we first introduce the mathematical notations employed in this study, followed by the proposed framework, omicsGAN, for generating synthetic omics data for disease outcome prediction using multi-omics data. The framework can take any two omics data with biological relations between each other as input. In this section, we used miRNA, mRNA, and miRNA-mRNA interaction network for illustrative purposes. We then discuss the evaluation metrics and introduce two evaluation methods; a classification model and a penalized Cox regression model that use the synthetic data for disease phenotype prediction and patient survival prediction, respectively.

### Overview of the framework

For the multi-omics data analysis, using extra omics data as an independent feature set provides additional information for downstream analysis. However, different omics profiles are often linked with each other through a complex biological interaction network. Our proposed framework, omicsGAN, can capture the information from this inter-omics network and integrate it with the omics datasets through a generative adversarial network to update them iteratively. After successful training of the network, it will generate new feature sets corresponding to each omics data that contain information from both modality and their interaction network. In this section, the framework is introduced on mRNA and miRNA expression datasets; however, this framework can work with any two omics data that are related to each other, given that their interactions are biologically meaningful. mRNA and miRNA expression are correlated to disease phenotype, although, the bipartite interaction network between them can be leveraged to increase the correlation by incorporating miRNA regulation on mRNA translation. mRNAs directly influence phenotype by translating into proteins that control all physiological activities in a cell; however, miRNA binds to mRNA and regulates its translation into protein, thus indirectly controls the phenotype. From a biological point of view, knowing the expression of a miRNA does not provide enough information without knowing the mRNAs that it targets. For an accurate and realistic down-stream analysis, realizing the interaction between omics data into calculation is crucial as well as challenging for the researchers.

The notations to define the proposed model, omicsGAN, are summarized in Table 1. Let ***N*** be the adjacency matrix of miRNA-mRNA interaction network and the dimension of the network is *p* ×*m*, where *p* is the number of miRNAs and *m* is the number of mRNAs. The dimensions of the mRNA (***X***) and miRNA (***Y***) expression data are *m*×*n* and *p*×*n* respectively, with *n* being the number of samples. Updated (synthetic) 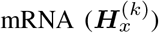 and 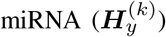 where *k* ∈ {1, 2, 3, …., *K*}, will correspond to the dimension of the input mRNA and miRNA expression datasets respectively and *K* is the total number of updates in omicsGAN.

**Table 1.**
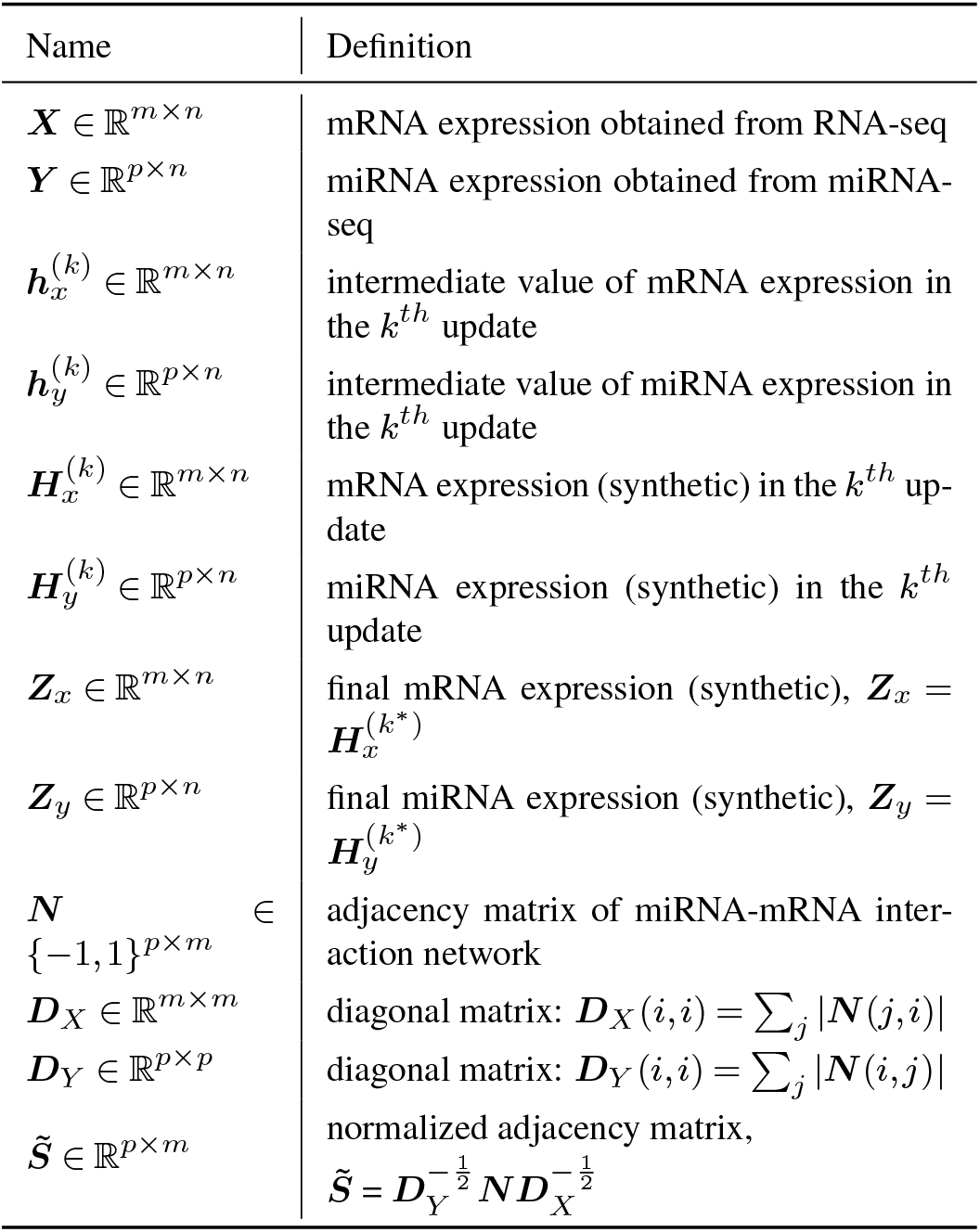
Notations for omicsGAN.

In this study, we predict disease outcome using two omics data and the interaction network between them as illustrated in Figure 1(a). The framework takes mRNA (***X***), miRNA (***Y***), and normalized interaction network 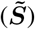 as input and iteratively updates them to find two new feature sets that incorporates information from both omics data and their biological interactions, where 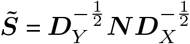. ***D***_*X*_ and ***D***_*Y*_ are two diagonal matrices with ***D***_*X*_ (*i, i*) =Σ_*j*_ |***N*** (*j, i*)| and ***D***_*Y*_ (*i, i*) =Σ_*j*_ |***N*** (*i, j*)|.. A classification model is then applied on the new feature sets to predict the disease phenotype. Figures 1(b) and (c) illustrate the frameworks for the first update (*k* = 1) of the mRNA and miRNA datasets respectively. Each box in Figure 1(a) represents *k*^*th*^ update which contains two Wasserstein GANs (wGANs) (22) for two omics data. After the wGANs are successfully trained, each generator generates a synthetic data which will be alike the input omics dataset and considered as the updated omics data from that box. For each update, an intermediate value for miRNA expression is first generated from the generator using mRNA expression and normalized adjacency matrix representing the interaction network. An intermediate value for the mRNA is also found in a similar procedure:

**Fig. 1.**
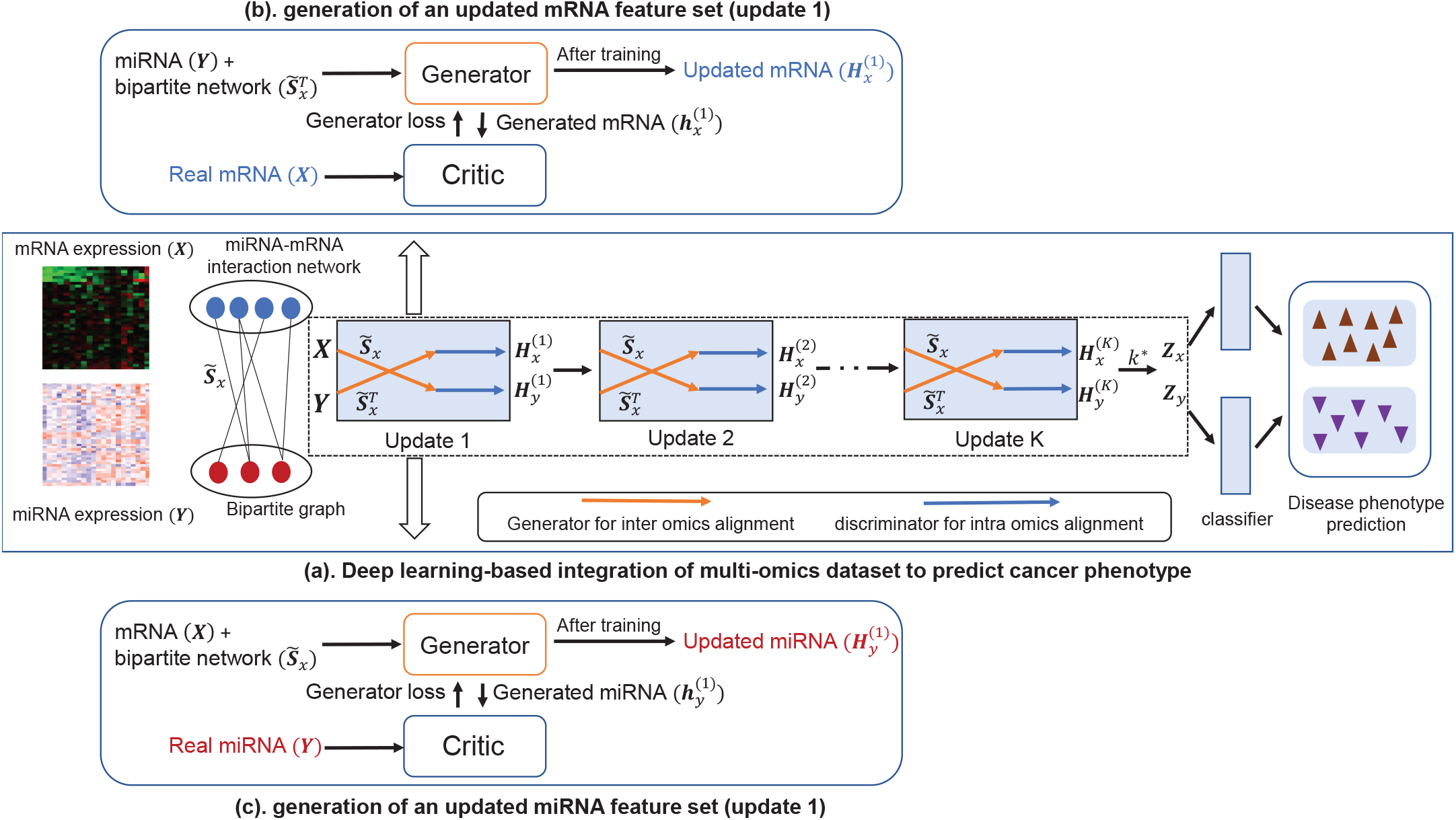
(a) An illustration of the proposed generative adversarial framework (omicsGAN). Two omics datasets are updated once in each box through an adversarial game between the generator (marked by orange line) and critic (marked by blue line). Generator and critic are trained for each omics data independently and the updated datasets are applied for disease phenotype prediction. **(b) Update of mRNA feature set**. Generator uses miRNA expression data and miRNA-mRNA bipartite network to synthesize an mRNA expression data. Both synthetic and input mRNA expression data are passed through a critic that tries to differentiate the real and synthetic data. **(c) Update of miRNA feature set**. Generator uses mRNA expression data and miRNA-mRNA bipartite network to synthesize an miRNA expression data. Both synthetic and input miRNA expression data are passed through a critic that tries to differentiate the real and synthetic data.

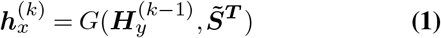

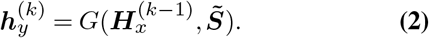

This mRNA (or miRNA) intermediate value 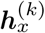 contains information from miRNA (mRNA) in the last update 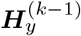 and interaction network 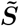 but has no relation with the mRNA (miRNA) expression value 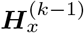 in the last update. The intermediate mRNA (or miRNA) expression value 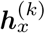 along with the input mRNA (miRNA) expression value 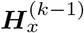 are then passed through a critic to ensure they are similar to each other:

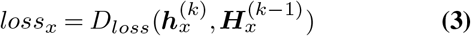

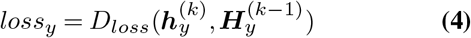

*D*_*loss*_ is the critic loss between the intermediate value and the input value. After training by minimizing the critic loss, the updated mRNA and miRNA dataset 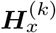 and 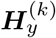 are learned respectively. This step force the distribution of 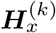 (or 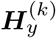) towards the distribution of 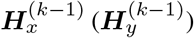. The boxes (updates) in Figure 1(a) are arranged in a cascaded structure where each box is trained separately. Once we have trained and got updates 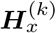 and 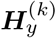 from box *k*, it is used as input in the following (*k*+1)^*th*^ box. 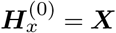 and 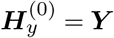 are the input to the first layer (box) and after the *K*^*th*^ update, 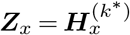 and 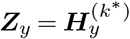 are our final synthetic datasets which are used for the disease phenotype prediction, where *k*^*\**^ is the update that gives best prediction result on a separated validation set of samples.

### A. Generative adversarial network model

Generative adversarial network (GAN) models are a class of unsupervised learning task that automatically discovers and learns patterns and distribution in input data in a way that the models can be used to generate new examples that plausibly could have been drawn from the original dataset. It has been widely used in image generation technologies (23, 24). With some appropriately placed conditions, it can also be used in computational biology to synthesize omics data. In general, GANs use random noise to generate synthetic dataset by requiring the distribution of the random noise towards the distribution of the original data. It does not have to retain information from the random noise; rather, try to make the noise as close to the original data as possible in terms of distribution. In multi-omics study, we can introduce a stream of information from one omics data in place of random noise and incentivize the GAN to retain information from this stream by using appropriate hyperparameters as well as forcing the distribution towards a second omics data. This will ensure the integration of information from both omics data in the generated samples. We can also fuse the interaction network in the model through the generator following the works of (25).

Our proposed pipeline has two separate wGANs for two omics data to update them into a new representation. Generators in each wGAN are four layers fully connected neural network that generates a dataset based on one omics data and the normalized adjacency matrix following the equations:

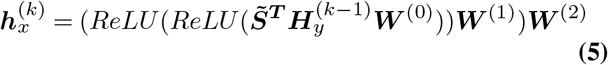

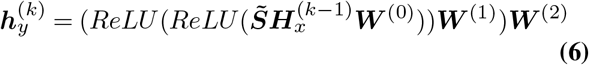

where ***W*** ^***l***^ is the weight matrix in *l*^*th*^ layer and rectified linear unit (ReLU) is the activation function. A fully connected neural network is then trained as a critic to assign values to the obtained intermediate representation 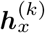 and input dataset 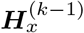. The critic is trained five times for one training of the generator. Objective function for training the critic is:

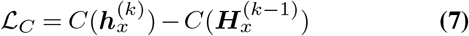

where *C* stands for the critic. Critic assigns larger values to the real samples 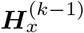 and smaller values to the synthetic ones 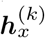, thus trained by minimizing equation (7). On the other hand, generator tries to produce synthetic data that will fool the critic into thinking it as real. Objective function for training the generator is:

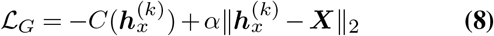

where *α* is a coefficient to control the weight put on the two terms of the equation. For a successful training, generator has to produce data 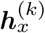 realistic enough that will be assigned a larger value by the critic; therefore, it is trained by minimizing equation (8). An *L*_2_-norm is added to further steer the updated dataset towards the original mRNA expression and preserve the feature characteristics. 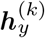 and 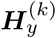 for miRNA update is derived using analogous equations.

### B. Evaluation methods. Classification model

We designed cancer outcome classification tasks with the assumption that better quality of the synthetic datasets will lead to better signatures for disease phenotype prediction compared to the original omics data. Support vector machine (SVM) with linear kernel is implemented as a classifier for all experiments. The datasets are divided into a ratio of 60%, 20%, 20% as numbers of training, validation, and test samples respectively. This model was implemented via Python package sklearn.svm (SVC).

#### Survival prediction model

A Cox proportional hazards model with Elastic Net penalty (26) is applied to study the correlation between patient’s overall survival and omics profiles. The Elastic Net penalty uses a weighted combination of the *L*_1_-norm and *L*_2_-norm penalties by maximizing the following log-likelihood function,

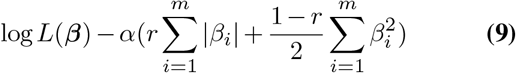

where *L*(*β*) is the partial likelihood of the Cox model, *α* ≥0 is a hyper-parameter that controls the amount of shrinkage, *r* ∈ [0, 1] is the relative weight of the *L*_1_-norm and *L*_2_-norm penalties, and *β*_*i*_(*I* ∈ [1, *m*]) represents the coefficient for the *i*^*th*^ genomic feature in the omics data. The omics data is randomly splitted into training (80%) and test (20%) sets. Five-fold cross validation is performed on training data to tune the hyper-parameter *α*. The high risk group and low risk group are determined by the prognostic index (***P I***) on the independent test set. The ***P I*** is the linear component of the Cox model, ***P I*** = *β*^*T*^ ***X***_*test*_, where ***X***_*test*_ is the omics profile of the test set, and its risk coefficient was estimated from the Cox model fitted on the training set. The high risk and low risk groups are generated for Kaplan-Meier survival plot by splitting the ordered ***P I*** with equal number of samples in each group in the test set. Python package *scikit survival* (27) is applied to implement Cox proportional hazards model with elastic net, and *lifelines* (28) is used for Kaplan-Meier plotting.

## Experiments

We performed experiments on The Cancer Genome Atlas (TCGA) datasets to evaluate the performance of omicsGAN with two different interaction networks (e.g., miRNA-mRNA interaction network and transcription factor (TF)-gene interaction network). In this section, we first describe the datasets and two interaction networks used in experiments. Next we introduce the experimental setup where we explain how to run our proposed model on TCGA data and generate synthetic omics datasets. Lastly, we performed three experiments to evaluate the performance of omicsGAN and the quality of its generated synthetic data: (1) comparing cancer outcome prediction power of the real and synthetic datasets.

The comparison was conducted in two ways: classifying clinical variables of cancer patients and number of significant features identified in each dataset; (2) exploring the impact of an accurate interaction network on the prediction power of synthetic datasets; (3) comparing the cancer patient’s overall survival prediction using real and synthetic datasets.

### C. Dataset and networks

The proposed framework, omicsGAN, was tested on The Cancer Genome Atlas (TCGA) breast invasive carcinoma (BRCA), lung adenocarcinoma (LUAD), and ovarian serous cystadenocarcinoma (OV) datasets (29–31). The RNA-seq mRNA expression and miRNA expression datasets of each cancer type were downloaded from UCSC Xena Hub (32). For the mRNA expression, the *log*2(*x* + 1) transformed RSEM normalized count with 20,531 genes was used and for the miRNA expression, the *log*2(*x* + 1) transformed RPM value with 2,166 miRNAs was used in this study. The clinical information of the three cancer studies was downloaded from cBioPortal (33). In breast cancer study, we classify the cancer patients based on estrogen receptor (ER+ vs ER-) and triple negative (TN+ vs TN-) status. Triple negative breast cancer patients test negative for all three receptors that are commonly found in breast cancer: estrogen receptors, progesterone receptors, and excess HER2 protein. For lung cancer and ovarian cancer studies, we classify the patients based on their survival time.

The miRNA-mRNA interaction network was obtained from TargetScanHuman (34). TargetScanHuman reports effective miRNA-mRNA interactions with context++ model, thereby providing valuable gene-regulatory networks with the miRNA involved. miRNA can bind to mRNA to cause more rapid degradation of the mRNA molecule, therefore reducing the amount of protein translated from that mRNA. A modified adjacency matrix represented the interaction network, where each interaction was valued as -1 to imitate that miRNA negatively regulates the expression of the targeted mRNA and no interaction was valued as 1. The miRNA-mRNA bipartite network contained 163,568 interactions in total. The TF-gene interaction network was downloaded from RegNetwork (35). The genes present in both lists of TFs and target genes were removed from the list of target genes. The modified bipartite interaction network contained sets of 1053 and 2859 non overlapping genes representing transcription factors and their target genes respectively with 8170 total interactions between them.

### D. Running omicsGAN on the TCGA datasets

To evaluate the proposed generative model on the TCGA omics datasets, we first updated the mRNA and miRNA (or TF and their target gene) expression profiles 5 times (*K* = 5). The generator and critic are fully connected neural networks with two hidden layers for the generator and one for the critic. The generator hidden layers have 512 and 768 neurons respectively whereas the critic hidden layers have 256 neurons. In both generator and critic, the activation function of the hidden layers is ReLU and the output layer is linear. Moreover, hidden layers in critic have dropout with a probability of 0.3. RMSprop optimizer was applied to train both the generator and the critic. Hyperparameters were selected through grid search and details of the hyperparameters used in this study are listed in Table 2. In Table 2, Omics 1 is the mRNA/gene expression data for both interaction networks, Omics 2 is miRNA expression in miRNA-mRNA interaction network and TF in TF-gene interaction network. The learning rate was chosen from {1e-8, 1e-7, 5e-7, 1e-6, 5e-6, 1e-5, 5e-5, 1e-4, 5e-4, 1e-3, 5e-3, 1e-2} and the candidates for the coefficient *α* were {1e-5, 1e-4, 1e-3, 1e-2, 0.1, 1, 10}. For batch size, we selected among the options {16, 32, 64, 128, 256}, and no mini batch. The validation set described in the Method section were employed for tuning all hyperparameters. All updated mRNA and miRNA (or gene and TF) datasets (*k* = 1, 2, ..., 5) are sequentially fed into the classifier. The support vector machine based classifier described in the Method section was used for classification in all experiments. In the classifier, the dataset was divided into five folds with three folds for training, one fold for validation (parameter tuning and synthetic data update selection), and one fold for testing. We repeated the five-fold splitting 50 times on each dataset. The updated mRNA/gene expression (*k*^*\**^) with the highest AUC score for validation samples was selected as the final synthetic mRNA/gene expression output from the model and similarly the updated miRNA/TF expression with the highest AUC score for validation samples was selected as the final synthetic miRNA/TF expression output. Figure 2 illustrates the process of selecting the final synthetic mRNA and miRNA datasets from all available updates for TCGA breast cancer patients outcome prediction. *k* = 1 gives the best validation AUC for synthetic mRNA expression whereas *k* = 2 gives the best validation AUC for synthetic miRNA expression. Therefore, mRNA update 1 and miRNA update 2 are used for predicting the test samples and the corresponding results are reported in this study. One synthetic data is generated for breast cancer ER and TN status prediction based on the average validation AUC of the two clinical variables.

**Table 2.** Hyperparameters in omicsGAN used in the study.

| Hyperparameter | miRNA-mRNA |  |  | TF-gene |
| --- | --- | --- | --- | --- |
|  | BRCA | LUAD | OV | LUAD |
| Omics 1 generator learning rate | 5e-6 | 5e-6 | 5e-6 | 5e-6 |
| Omics 1 critic learning rate | 5e-5 | 5e-5 | 5e-5 | 5e-5 |
| Omics 1 $L_2$-norm coefficient ($\alpha$) | 0.01 | 0.01 | 0.1 | 0.0001 |
| Omics 2 generator learning rate | 5e-6 | 5e-6 | 5e-6 | 5e-6 |
| Omics 2 critic learning rate | 5e-5 | 5e-5 | 5e-5 | 5e-5 |
| Omics 2 $L_2$-norm coefficient ($\alpha$) | 0.001 | 0.001 | 0.001 | 0.001 |

**Fig. 2.**
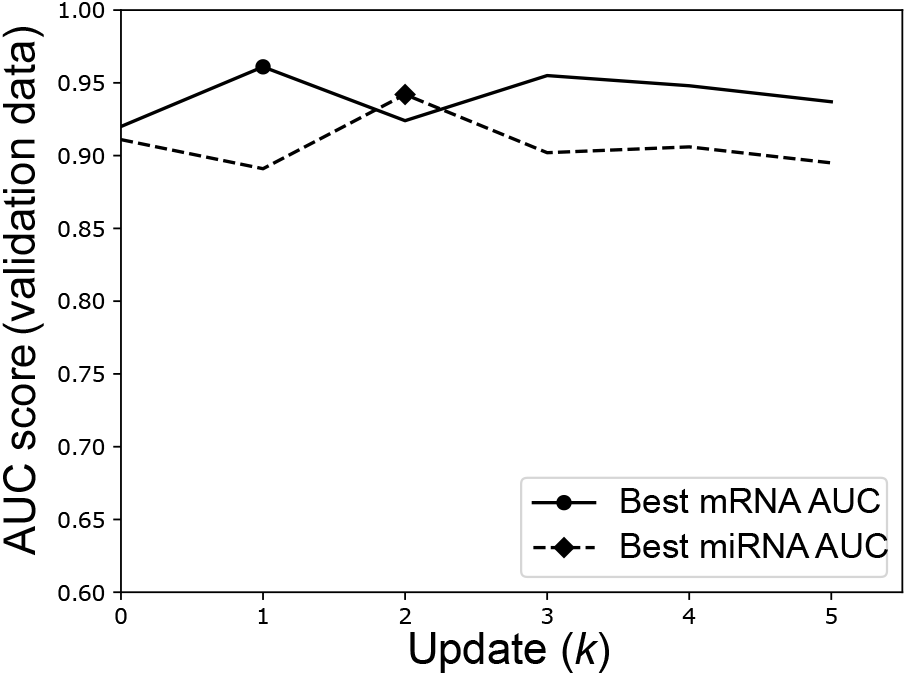
Prediction results of triple negative (TN) status on TCGA breast cancer patients using validation samples. AUC of the prediction results using validation samples of synthetic mRNA and miRNA for *k* = [1, 2, 3, 4, 5]. Update *k*^*\**^ with the best validation AUC is selected as the final synthetic data for each omics profile.

### E. Integration of mRNA and miRNA expression

We generate the synthetic mRNA and miRNA datasets by integrating the two omcis profiles and their interaction network and assess the quality of the synthetic data through three experiments.

#### E.1. omicsGAN improved cancer outcome prediction

To evaluate the quality of the synthetic datasets generated by omicsGAN, we designed cancer outcome prediction and significant predictive signature identification tasks on the TCGA breast cancer, lung cancer, and ovarian cancer datasets under the assumptions: (1) The synthetic datasets learned in omics-GAN consider the expressions in both mRNA and miRNA profiles and the biological interactions between them. So they will provide better predictive signatures compared to mRNA and miRNA expressions. (2) The better predictive signatures will improve the disease phenotype prediction.

We ran the classifier with above mentioned five-fold splitting 50 times to select the best synthetic data among the 5 updates based on validation samples and classify the test samples using the selected synthetic data. The average AUC scores of 50 splittings are reported in Table 3. There are 185 Estrogen Receptor positive (ER+) and 54 ER negative (ER-) samples, 46 triple negative positive (TN+) and 193 TN negative (TN-) samples in the breast cancer dataset, 95 cancer patients below the survival time cutoff (*<* 25 months) and 64 above the cutoff (*>* 50 months) in the lung cancer dataset as well as 61 cancer patients below the survival time cutoff (*<* 25 months) and 77 above the cutoff (*>* 50 months) in the ovarian cancer dataset. Table 3 illustrates that the synthetic mRNA and miRNA expression generated by omicsGAN achieved better average classification results than original mRNA and miRNA expression for phenotype predictions across all three cancer types. We also add the baseline where we perform the classification with concatenated miRNA and mRNA expression to see whether addition of more omics data is the reason for the improvement. We can see that concatenated data has similar or better prediction ability compared to the original mRNA and miRNA expression dataset; however, synthetic dataset from omicsGAN always outperforms the concatenated data by a significant margin. This signifies that even though the addition of more omics data improves the outcome prediction performance, omicsGAN relies on the interaction network to generate synthetic data with better predictive signal.

**Table 3.**
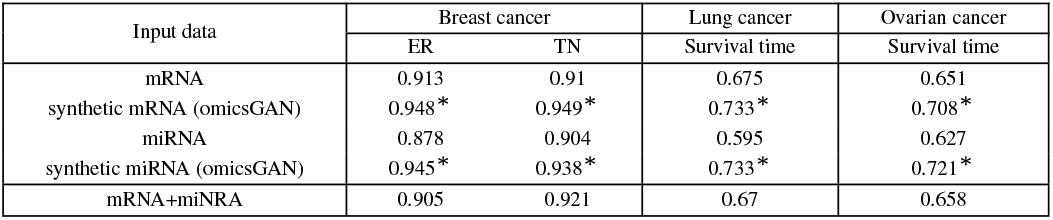
The classification performance on TCGA breast cancer, lung cancer, and ovarian cancer datasets. Average AUC scores of classify cancer patients clinical variables on the synthetic mRNA, miRNA datasets generated from omicsGAN and the original mRNA, miRNA expression datasets. *\** The difference between the results on the original expression data and the synthetic data is statistically significant (*p*-value < 0.001).

| Input data | Breast cancer |  | Lung cancer | Ovarian cancer |
| --- | --- | --- | --- | --- |
|  | ER | TN | Survival time | Survival time |
| mRNA | 0.913 | 0.91 | 0.675 | 0.651 |
| synthetic mRNA (omicsGAN) | 0.948* | 0.949* | 0.733* | 0.708* |
| miRNA | 0.878 | 0.904 | 0.595 | 0.627 |
| synthetic miRNA (omicsGAN) | 0.945* | 0.938* | 0.733* | 0.721* |
| mRNA+miRNA | 0.905 | 0.921 | 0.67 | 0.658 |

We also evaluated the quality of the original and synthetic datasets by comparing the number of significant features identified in each of them. We performed Student’s *t*-test on the expression datasets with different clinical variables. The number of features with a *p*-value smaller than 0.001 in each dataset except miRNA expression for lung cancer patients are presented in Table 4. *p*-value cutoff of 0.05 is set for miRNA expression for lung cancer patients as no feature had a *p*-value smaller than 0.001 in either the real miRNA expression or the synthetic one. We can see an increased number of significant features in synthetic mRNA compared to the original one for all three cancer types. Synthetic miRNA on the other hand has more significant features for breast cancer and lung cancer, but less for ovarian cancer compared to the original miRNA expression datasets. Therefore, omicsGAN enriches the features of synthetic datasets with better predictive signatures that results into improved cancer outcome prediction.

#### E.2. Impact of interaction network on cancer outcome prediction

miRNA expression provides additional predictive signals for cancer outcome prediction on top of the mRNA expression; therefore, integrating them into a new feature set will contain more information compared to mRNA and miRNA expression individually. Table 3 and 4 already illustrates the ability of omicsGAN to improve the cancer outcome prediction performance. However, we hypothesized that omicsGAN harnesses the information of biological interaction between two omics layers from multi-omics interaction network to generate the synthetic datasets with better predictive signals. Hence, we want to investigate whether the improvement in performance is because of the additional omics data or the model can exploit the interaction network for data integration. We design an experiment to explore the effects of the interaction network on synthetic omics data and their predictive performance where we ran the framework 10 times with same settings and input ***X*** (mRNA expression), ***Y*** (miRNA expression) as before but a different interaction network on TCGA lung cancer datasets. We replaced the true network with 10 different randomized networks with same density as the true one. The prediction results for synthetic mRNA and miRNA expression using true and random networks are shown as boxplots in Figures 3 and 4 respectively. Prediction results using original mRNA/miRNA expression, synthetic mRNA/miRNA expression generated using the true network, and synthetic mRNA/miRNA expression generated using random network are plotted in each figure. The first two boxplots display the same results for lung cancer outcome prediction as shown in Table 3. 50 dots in each of these two boxplots represent the AUC corresponding to 50 random splittings. The third boxplot illustrates the results using 10 random networks, each with 50 splittings. The statistics (mean, median, and standard deviation) of the prediction performance of the splittings are shown above each boxplot. In Figures 3 and 4, we see a reduction in performance of synthetic mRNA/miRNA expression generated using a random interaction network compared to the one generated using the true interaction network. This signifies the importance of the interaction network in phenotype prediction and the capability of our framework to capture the information within the network.

**Table 4.** Number of significant features. Number of significant features between synthetic mRNA, miRNA generated by omicsGAN and the original mRNA, miRNA expression on breast cancer, lung cancer, and ovarian cancer datasets.

| Input data | Breast cancer |  | Lung cancer | Ovarian cancer |
| --- | --- | --- | --- | --- |
|  | ER | TN | Survival time | Survival time |
| mRNA | 4144 | 3893 | 227 | 133 |
| synthetic mRNA (omicsGAN) | 4566 | 4241 | 372 | 142 |
| miRNA | 91 | 91 | 23 | 20 |
| synthetic miRNA (omicsGAN) | 136 | 127 | 58 | 12 |

**Fig. 3.**
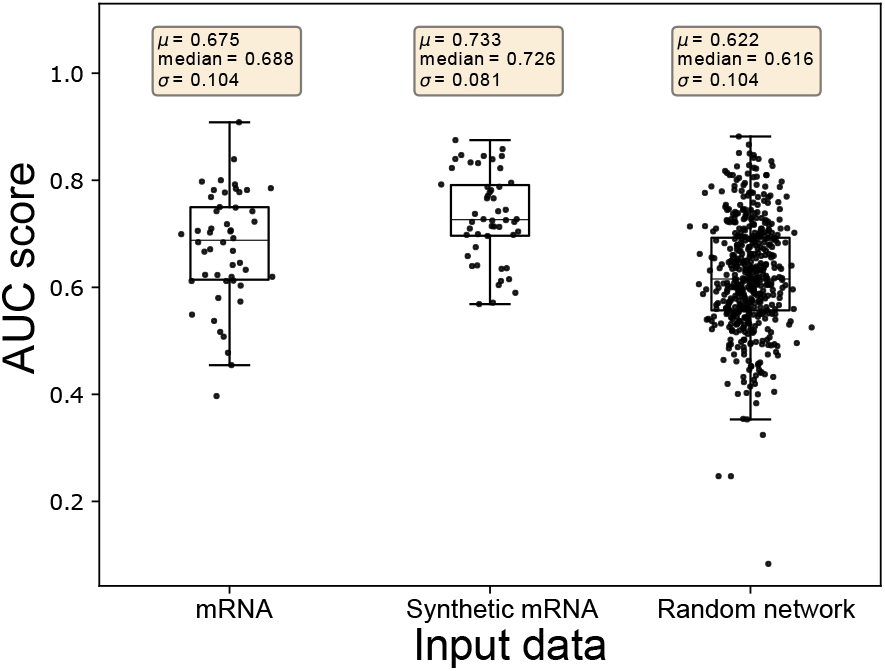
Prediction results of the survival time on TCGA lung cancer patients using original and synthetic mRNA expression. Prediction results using original mRNA expression, synthetic mRNA expression generated using true interaction network, and synthetic mRNA expression generated using random interaction network are plotted respectively. Each dot represents the AUC score from one splitting. Statistics (mean, median,and standard deviation) of the prediction performance of the 50 splittings are shown above each boxplot.

**Fig. 4.**
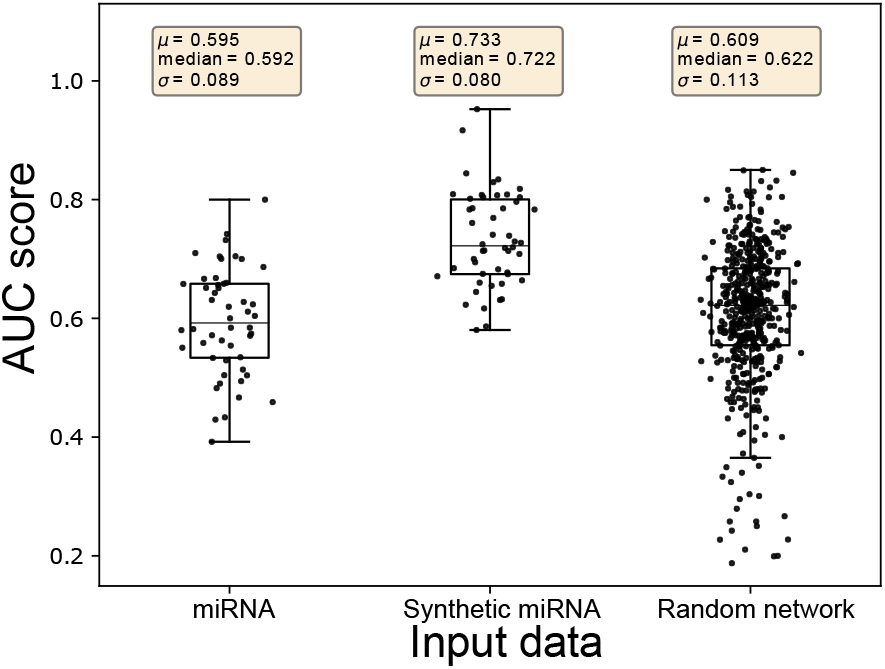
Prediction results of the survival time on TCGA lung cancer patients using original and synthetic miRNA expression. Prediction results using original miRNA expression, synthetic miRNA expression generated using true interaction network, and synthetic miRNA expression generated using random interaction network are plotted respectively. Each dot represents the AUC score from one splitting. Statistics (mean, median,and standard deviation) of the prediction performance of the 50 splittings are shown above each boxplot.

#### E.3. omicsGAN improved survival prediction

To further investigate the quality of the synthetic mRNA and miRNA expression data produced by omicsGAN, the patient’s overall survival was predicted on breast cancer, lung cancer, and ovarian cancer datasets. The Cox proportional hazards model with elastic net penalty as described in section B evaluates the correlation between patient’s overall survival and genomic features, i.e., the original mRNA, miRNA expressions and the synthetic mRNA, miRNA expressions in this study. The relative weight *r* in equation 9 was set to be 0.5 to combine the subset selection property of the *L*_1_-norm with the regularization strength of the *L*_2_-norm. 80% of the patient samples were applied to train the model and the performance was tested on 20% test samples. The low and high risk groups on the independent test set were generated based on the prognostic index (***P I***) as mentioned in section B. The survival predictions were visualized by Kaplan-Meier plots and compared by the log-rank test *p*-values. The Kaplan-Meier plots in Figure 5 and 6 exemplify the improved patient survival predictions on lung cancer using the synthetic mRNA, miRNA expressions generated by omicsGAN compared to the original mRNA, miRNA expressions. The log-rank test *p*-values clearly demonstrate a strong additional prognostic power of the synthetic omics profiles beyond the original signatures. Similar observations are identified on breast and ovarian cancer patient samples (Supplementary Figure 1-4).

**Fig. 5.**
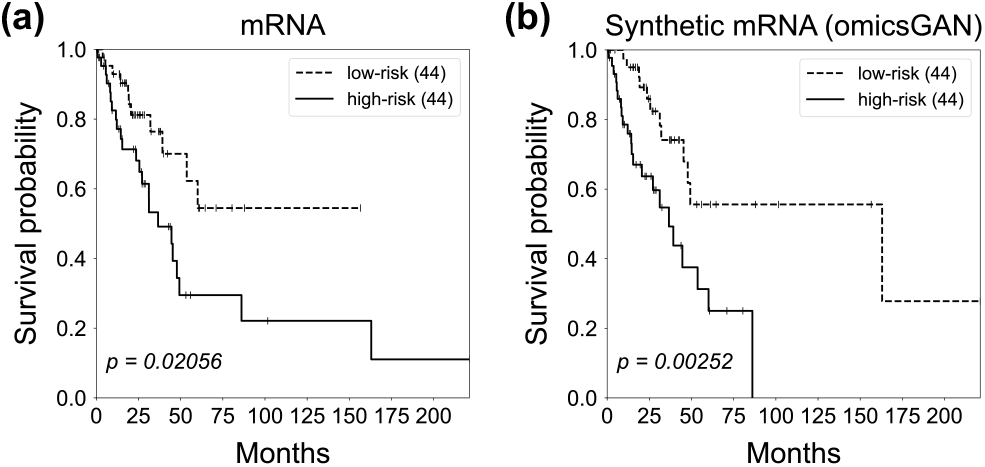
Survival prediction on lung cancer patients with mRNA profiles. Kaplan-Meier survival plots for high (solid line) and low (dashed line) risk groups generated by (a) original mRNA, (b) synthetic mRNA expression data on lung cancer patients. The number in the parenthesis indicates the number of samples in low or high risk group. The *p*-value is calculated by the log-rank test to compare the overall survival of two groups of cancer patients.

**Fig. 6.**
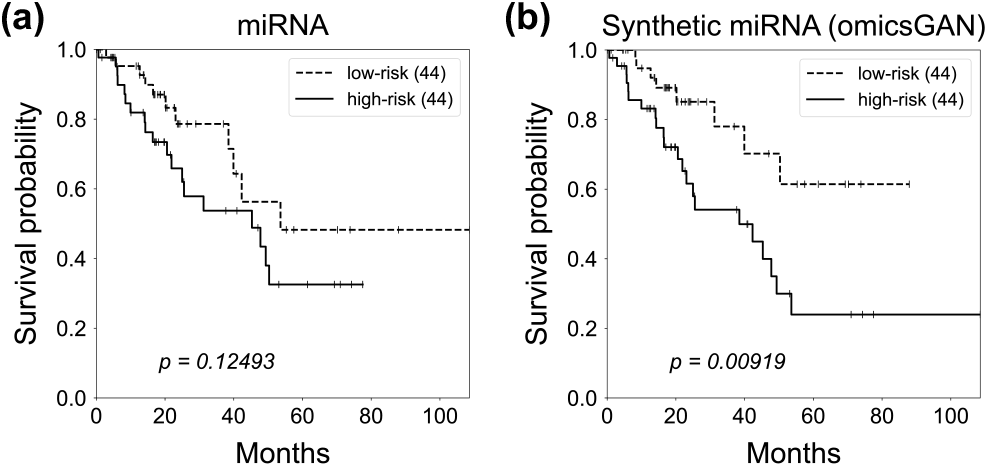
Survival prediction on lung cancer patients with miRNA profiles. Kaplan-Meier survival plots for high (solid line) and low (dashed line) risk groups generated by (a) original miRNA, (b) synthetic miRNA expression data on lung cancer patients. The number in the parenthesis indicates the number of samples in low or high risk group. The *p*-value is calculated by the log-rank test to compare the overall survival of two groups of cancer patients.

#### F. Integration of transcription factor and gene expression

The experiments above shows the ability of omicsGAN to generate synthetic data with better predictive power by harnessing the information from miRNA-mRNA interaction network. Here, we design another experiment using transcription factor (TF)-gene interaction network to evaluate whether omicsGAN can show similar improvement in integrating other omics data and their interaction network. We performed the lung cancer phenotype prediction based on the same classification setup as described in section E.1 on TFs and their target gene expression datasets. The average AUC scores of 50 splittings are reported in Table 5. Both the synthetic TF and target gene expression performed better in classifying the lung cancer patients based on their survival time than the original TF, gene expression, and concatenated TF and gene expression. These findings signify that our proposed framework can work with varying set multi-omics data.

**Table 5.** The classification performance on TCGA lung cancer dataset. Average AUC scores of classification performance between synthetic gene, TF generated from omicsGAN and the original gene, TF expression on lung cancer datasets.

| Input data | Lung cancer |
| --- | --- |
| gene | 0.645 |
| synthetic gene | 0.727* |
| TF | 0.656 |
| synthetic TF | 0.743* |
| gene+TF | 0.682 |

## Discussion

Disease phenotype prediction plays a key role in the fight against heterogeneous diseases like cancer. Multi-omics data powered by next generation sequencing technologies has transformed the field of phenotype prediction by providing a broader view of the molecular profiles. Non-redundant predictive signals from multi-omics data make it crucial to develop an efficient and effective framework for multi-omics data integration. However, integrating them as an independent set of features is inadequate as multi-omics data generated for the same set of samples often have an interactive relation among them. Incorporating the interaction network into the analysis will set a flow of information from one omics data to another like the flow within different omics layers in a cell. In most studies, these inter-omics relations are neglected and it is inefficient to predict phenotype using integrated multi-omics data without considering the interactions. Therefore, the integrating of the bipartite interaction network with multi-omics data can result in improved disease phenotype prediction and designing frameworks capable of such integration is gaining importance.

Synthetic data generated from our proposed framework, omicsGAN, shows improvement in prediction performance which illustrates the capability of the model to successfully retain information from multiple omics data and establish a link between them. All synthetic datasets generated in this study with two interaction networks (i.e., miRNA-mRNA and TF-gene) perform better in cancer outcome prediction compared to the original expression datasets; however, the same model using a random interaction network with same density does not perform as good as the synthetic datasets obtained through true network. It signifies that omicsGAN does not fuse information from the two omics data directly; rather functionally incorporate the interaction network into the integration. Synthetic miRNA expression using random interaction network works better than the original miRNA expression (Figure 4) but synthetic mRNA using random interaction network does not perform better than original mRNA expression (Figure 3). The reason is, without the true interaction network, omicsGAN can still integrate information from the two omics data to generate synthetic datasets. In that case, the performance of one synthetic data will depend on the additional information received from the other omics data. Synthetic miRNA receives information from mRNA expression, which is significantly better in lung cancer outcome prediction compared to miRNA and thus improves the performance of synthetic miRNA. Synthetic mRNA on the other hand receives information from miRNA that is worse at prediction compared to mRNA and thus results in a decreased performance. An *L*_2_-norm is added in equation (8) to ensure the similarity between the updated and original omics data expression; thus allowing the synthetic data to retain feature space properties of the original omics data.

The framework presents an innovative way for multi-omics data integration incorporating their biological interaction. A larger comprehensive study involving more cancer types can draw a better picture of the improvements in phenotype prediction. Although our study was focused on miRNA-mRNA interaction and TF-gene interaction, the same technique can be extrapolated to any two omics data if their interaction network is biologically meaningful. However, to integrate two omics data with different range, distribution, and format (e.g., mutation and gene expression), an extra pre-processing step is necessary to make them compatible. There are existing studies with the pre-processing framework that works with such multi-omics data (36).

## Conclusion

Thanks to the rapid evolution of high-throughput technologies, abundant genotype data is accruing, which is expected to grow continuously in the era of precision medicine. Because of the complex interactive nature of omics layers, integration of multi-omics data to extract biologically meaningful information of clinical relevance is a challenging task. The promise of multi-omics analysis will remain unfulfilled unless we can functionally incorporate the inter-omics interaction network into the analysis. In this study, we introduced omicsGAN, a generative adversarial network model to effectively integrate the interaction network and the omics datasets into new synthetic data with better predictive signals. We observed that the synthetic data generated from omicsGAN has better discriminative power on cancer outcome classification and cancer patients survival prediction compared to the original omics datasets. Synthetic datasets also contain more significant features that result in better predictive performance. Additionally, we analyzed the effect of interaction network on the quality of synthetic data. Our results show that omicsGAN does not only gather information from two omics datasets; rather functionally incorporate their biological interaction into the integration. Using a random interaction network does not create a flow of information from one omics data to another as efficiently as the true network.

## Acknowledgements

The results are based upon data generated by The Cancer Genome Atlas established by the NCI and NHGRI. Information about TCGA and the investigators and institutions who constitute the TCGA research network can be found at http://cancergenome.nih.gov. The dbGaP accession number to the specific version of the TCGA dataset is phs000178.v8.p7.

## Bibliography

1. Paulina Krzyszczyk, Alison Acevedo, Erika J Davidoff, Lauren M Timmins, Ileana Marrero-Berrios, Misaal Patel, Corina White, Christopher Lowe, Joseph J Sherba, Clara Hartmanshenn, et al. The growing role of precision and personalized medicine for cancer treatment. Technology, 6(03n04):79–100, 2018.

2. Hugo Fitipaldi, Mark I McCarthy, Jose C Florez, and Paul W Franks. A global overview of precision medicine in type 2 diabetes. Diabetes, 67(10):1911–1922, 2018.

3. Chao Wang, Raghu Machiraju, and Kun Huang. Breast cancer patient stratification using a molecular regularized consensus clustering method. Methods, 67(3):304–312, 2014.

4. Sara Goodwin, John D McPherson, and W Richard McCombie. Coming of age: ten years of next-generation sequencing technologies. Nature Reviews Genetics, 17(6):333, 2016.

5. Indhupriya Subramanian, Srikant Verma, Shiva Kumar, Abhay Jere, and Krishanpal Anamika. Multi-omics data integration, interpretation, and its application. Bioinformatics and biology insights, 14:1177932219899051, 2020.

6. Bo Wang, Aziz M Mezlini, Feyyaz Demir, Marc Fiume, Zhuowen Tu, Michael Brudno, Benjamin Haibe-Kains, and Anna Goldenberg. Similarity network fusion for aggregating data types on a genomic scale. Nature methods, 11(3):333, 2014.

7. Marc Vidal, Michael E Cusick, and Albert-László Barabási. Interactome networks and human disease. Cell, 144(6):986–998, 2011.

8. Hsin-Sung Yeh, Wei Zhang, and Jeongsik Yong. Analyses of alternative polyadenylation: from old school biochemistry to high-throughput technologies. BMB reports, 50(4):201, 2017.

9. Julia K Nussbacher and Gene W Yeo. Systematic discovery of RNA binding proteins that regulate microRNA levels. Molecular cell, 69(6):1005–1016, 2018.

10. Matthias W Hentze, Alfredo Castello, Thomas Schwarzl, and Thomas Preiss. A brave new world of RNA-binding proteins. Nature Reviews Molecular Cell Biology, 19(5):327, 2018.

11. Nimrod Rappoport and Ron Shamir. NEMO: Cancer subtyping by integration of partial multi-omic data. Bioinformatics, 35(18):3348–3356, 2019.

12. Hung Nguyen, Sangam Shrestha, Sorin Draghici, and Tin Nguyen. PINSPlus: a tool for tumor subtype discovery in integrated genomic data. Bioinformatics, 35(16):2843–2846, 2019.

13. Ricard Argelaguet, Britta Velten, Damien Arnol, Sascha Dietrich, Thorsten Zenz, John C Marioni, Florian Buettner, Wolfgang Huber, and Oliver Stegle. Multi-Omics Factor Analysis—a framework for unsupervised integration of multi-omics data sets. Molecular systems biology, 14(6):e8124, 2018.

14. Tao Zhou, Kim-Han Thung, Xiaofeng Zhu, and Dinggang Shen. Effective feature learning and fusion of multimodality data using stage-wise deep neural network for dementia diagnosis. Human brain mapping, 40(3):1001–1016, 2019.

15. Hiromi WL Koh, Damian Fermin, Christine Vogel, Kwok Pui Choi, Rob M Ewing, and Hyungwon Choi. iOmicsPASS: network-based integration of multiomics data for predictive subnetwork discovery. NPJ systems biology and applications, 5(1):1–10, 2019.

16. Ian Goodfellow, Jean Pouget-Abadie, Mehdi Mirza, Bing Xu, David Warde-Farley, Sherjil Ozair, Aaron Courville, and Yoshua Bengio. Generative adversarial nets. Advances in neural information processing systems, 27:2672–2680, 2014.

17. Minseon Kim, Ilhwan Oh, and Jaegyoon Ahn. An improved method for prediction of cancer prognosis by network learning. Genes, 9(10):478, 2018.

18. Arsham Ghahramani, Fiona M Watt, and Nicholas M Luscombe. Generative adversarial networks simulate gene expression and predict perturbations in single cells. BioRxiv, page 262501, 2018.

19. Jinhee Park, Hyerin Kim, Jaekwang Kim, and Mookyung Cheon. A practical application of generative adversarial networks for RNA-seq analysis to predict the molecular progress of Alzheimer’s disease. PLoS computational biology, 16(7):e1008099, 2020.

20. Mojtaba Bahrami, Malosree Maitra, Corina Nagy, Gustavo Turecki, Hamid Rabiee, and Yue Li. Deep feature extraction of single-cell transcriptomes by generative adversarial network. bioRxiv, 2020.

21. Yungang Xu, Zhigang Zhang, Lei You, Jiajia Liu, Zhiwei Fan, and Xiaobo Zhou. scIGANs: single-cell RNA-seq imputation using generative adversarial networks. Nucleic acids research, 48(15):e85–e85, 2020.

22. Martin Arjovsky, Soumith Chintala, and Léon Bottou. Wasserstein gan. arXiv preprint arXiv:1701.07875, 2017.

23. Yunjey Choi, Minje Choi, Munyoung Kim, Jung-Woo Ha, Sunghun Kim, and Jaegul Choo. Stargan: Unified generative adversarial networks for multi-domain image-to-image translation. In Proceedings of the IEEE conference on computer vision and pattern recognition, pages 8789–8797, 2018.

24. Han Zhang, Tao Xu, Hongsheng Li, Shaoting Zhang, Xiaogang Wang, Xiaolei Huang, and Dimitris N Metaxas. Stackgan: Text to photo-realistic image synthesis with stacked generative adversarial networks. In Proceedings of the IEEE international conference on computer vision, pages 5907–5915, 2017.

25. Thomas N Kipf and Max Welling. Semi-supervised classification with graph convolutional networks. arXiv preprint 1609.02907, 2016.

26. Noah Simon, Jerome Friedman, Trevor Hastie, and Rob Tibshirani. Regularization paths for Cox’s proportional hazards model via coordinate descent. Journal of statistical software, 39 (5):1, 2011.

27. Sebastian Pölsterl. scikit-survival: A Library for Time-to-Event Analysis Built on Top of scikit-learn. Journal of Machine Learning Research, 21(212):1–6, 2020.

28. Cameron Davidson-Pilon, Jonas Kalderstam, Noah Jacobson, Sean Reed, Ben Kuhn, Paul Zivich, Mike Williamson, Abdeali JK, Deepyaman Datta, Andrew Fiore-Gartland, Alex Parij, Daniel WIlson, Gabriel Luis Moneda, Arturo Moncada-Torres, Kyle Stark, Harsh Gadgil, Jona Karthikeyan Singaravelan, Lilian Besson, Miguel Sancho Peña Steven Anton, Andreas Klintberg, GrowthJeff Javad Noorbakhsh, Matthew Begun, Ravin Kumar, Sean Hussey, Skipper Seabold, and Dave Golland. CamDavidsonPilon/lifelines: v0.25.8, January 2021.

29. Cancer Genome Atlas Network TCGA et al. Comprehensive molecular portraits of human breast tumours. Nature, 490(7418):61, 2012.

30. Cancer Genome Atlas Research Network TCGA et al. Comprehensive molecular profiling of lung adenocarcinoma. Nature, 511(7511):543, 2014.

31. Cancer Genome Atlas Research Network TCGA et al. Integrated genomic analyses of ovarian carcinoma. Nature, 474(7353):609, 2011.

32. Mary J Goldman, Brian Craft, Mim Hastie, Kristupas Repečka, Fran McDade, Akhil Kamath, Ayan Banerjee, Yunhai Luo, Dave Rogers, Angela N Brooks, et al. Visualizing and interpreting cancer genomics data via the Xena platform. Nature Biotechnology, pages 1–4, 2020.

33. Jianjiong Gao, Bülent Arman Aksoy, Ugur Dogrusoz, Gideon Dresdner, Benjamin Gross, S Onur Sumer, Yichao Sun, Anders Jacobsen, Rileen Sinha, Erik Larsson, et al. Integrative analysis of complex cancer genomics and clinical profiles using the cBioPortal. Science signaling, 6(269):pl1–pl1, 2013.

34. Vikram Agarwal, George W Bell, Jin-Wu Nam, and David P Bartel. Predicting effective microRNA target sites in mammalian mRNAs. elife, 4:e05005, 2015.

35. Zhi-Ping Liu, Canglin Wu, Hongyu Miao, and Hulin Wu. RegNetwork: an integrated database of transcriptional and post-transcriptional regulatory networks in human and mouse. Database, 2015, 2015.

36. Hossein Sharifi-Noghabi, Olga Zolotareva, Colin C Collins, and Martin Ester. MOLI: multiomics late integration with deep neural networks for drug response prediction. Bioinformatics, 35(14):i501–i509, 2019.

