## Supplementary Figure 1-4 for "Multi-omics Data Integration by Generative Adversarial Network"

### 1 Kaplan-Meier survival plots on breast and ovarian cancer

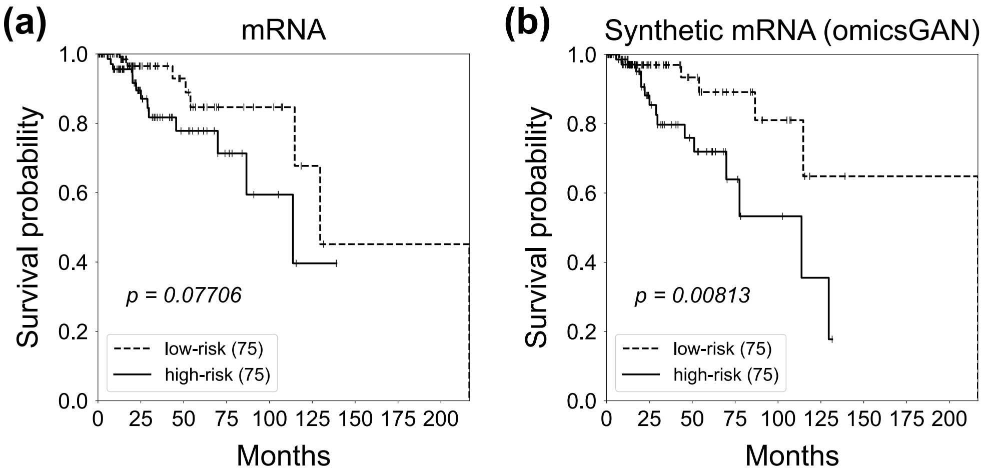


Figure 1: **Survival prediction on breast cancer patients with mRNA profiles.** Kaplan-Meier survival plots for high (solid line) and low (dashed line) risk groups generated by (a) original mRNA, (b) synthetic mRNA expression data on breast cancer patients. The number in the parenthesis indicates the number of samples in low or high risk group. The $p$-value is calculated by the log-rank test to compare the overall survival of two groups of cancer patients.

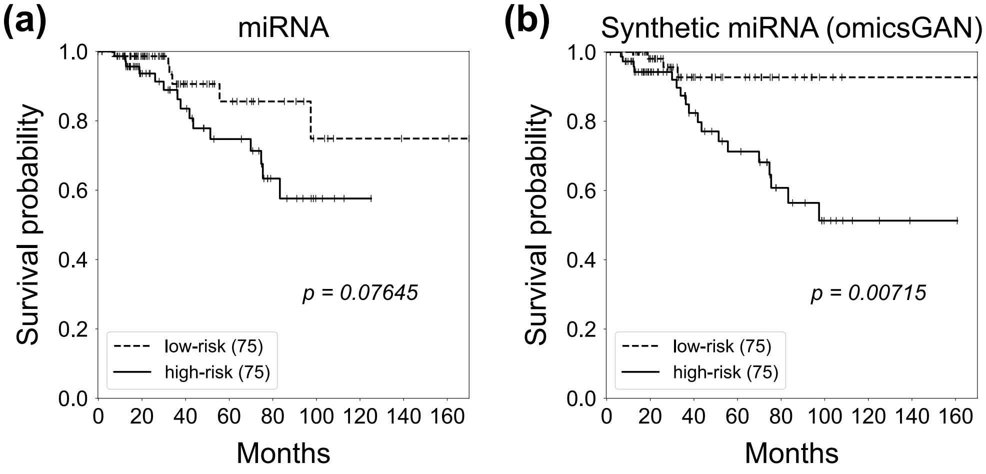


Figure 2: **Survival prediction on breast cancer patients with miRNA profiles.** Kaplan-Meier survival plots for high (solid line) and low (dashed line) risk groups generated by (a) original miRNA, (b) synthetic miRNA expression data on breast cancer patients. The number in the parenthesis indicates the number of samples in low or high risk group. The $p$-value is calculated by the log-rank test to compare the overall survival of two groups of cancer patients.

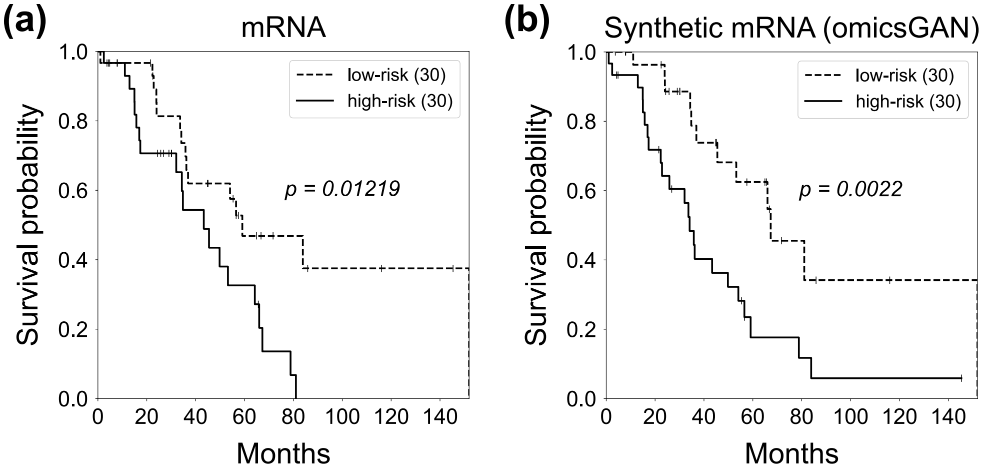


Figure 3: **Survival prediction on ovarian cancer patients with mRNA profiles.** Kaplan-Meier survival plots for high (solid line) and low (dashed line) risk groups generated by (a) original mRNA, (b) synthetic mRNA expression data on ovarian cancer patients. The number in the parenthesis indicates the number of samples in low or high risk group. The $p$-value is calculated by the log-rank test to compare the overall survival of two groups of cancer patients.

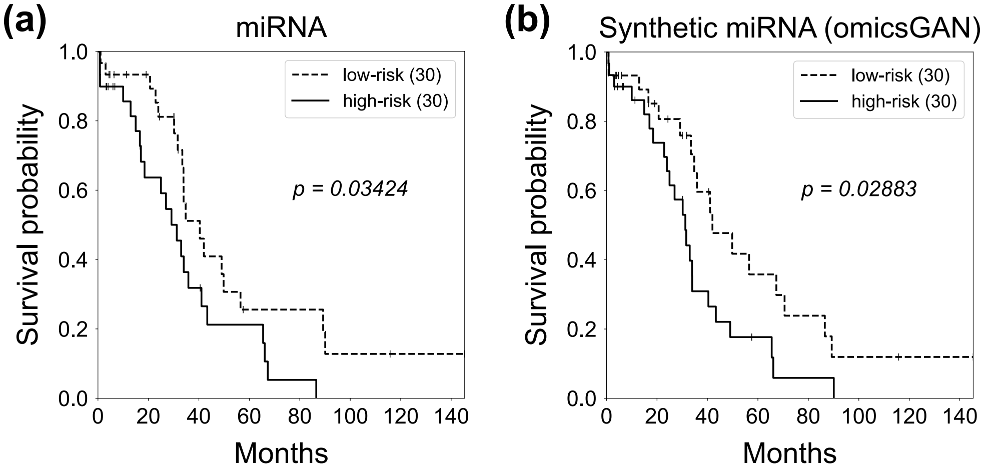


Figure 4: **Survival prediction on ovarian cancer patients with miRNA profiles.** Kaplan-Meier survival plots for high (solid line) and low (dashed line) risk groups generated by (a) original miRNA, (b) synthetic miRNA expression data on ovarian cancer patients. The number in the parenthesis indicates the number of samples in low or high risk group. The $p$-value is calculated by the log-rank test to compare the overall survival of two groups of cancer patients.
